# Optical flow analysis dissociates cortical selectivity for body movements and shapes

**DOI:** 10.64898/2026.02.26.708181

**Authors:** Baichen Li, Rufin Vogels, Giuseppe Marrazzo, Marta Poyo Solanas, Beatrice de Gelder

**Author notes:** Correspondence: Beatrice de Gelder < >.

## Abstract

Perceiving the movements of other living organisms is a fundamental ability across species. Although several cortical regions are selective for static body features, it is unclear whether these areas also encode motion cues independently of body shapes. Here, we isolated optical flow patterns from naturalistic body movements using synthesized videos that eliminate explicit body shape information. Behavioral results suggest that participants successfully recognized walking directions from the shape-free flow motion stimuli. Using ultra-high field fMRI, we found that three body-motion-related areas, fusiform gyrus (FG), lateral occipitotemporal cortex (LOTC), and posterior superior temporal sulcus (pSTS), were sensitive to both the amount and the spatiotemporal structure of flow motion. Critically, the flow motion effects were significantly correlated to voxel-wise body motion selectivity in FG and LOTC, but not to body shape selectivity in any of the ROIs. These findings identify flow motion as a distinct and behaviorally sufficient cue for body perception, which could dissociate the neural mechanism underlying motion and shape encoding in FG and LOTC.

## Introduction

Perceiving the movements of other living organisms is a fundamental survival skill shared across species. A large body of research has shown that human observers can readily recognize others’ intentions, actions, and emotions from brief whole-body movements (De Gelder, 2006; De Meijer, 1989; Roether et al., 2009; Wallbott, 1998). However, the neural mechanisms underlying body perception and motion perception remain largely disconnected. Research on visual motion analysis has primarily focused on area MT and the dorsal visual pathway (Newsome et al., 1990; Parker & Newsome, 1998; Rust et al., 2006), whereas studies on body perception have mainly been grounded in the ventral visual pathway associated with object recognition (Peelen & Downing, 2017; Ritchie et al., 2025). Distinct neural representations for body features have been reported in the lateral occipitotemporal cortex (LOTC) and fusiform gyrus (FG), specifically in the extrastriate body area (EBA; Downing et al., 2001; Downing et al., 2006; Jastorff & Orban, 2009; Taylor et al., 2007; Vangeneugden et al., 2014) and in the fusiform body area (FBA; Jastorff & Orban, 2009; Peelen & Downing, 2005; Taylor et al., 2007). Although these body-selective regions are widely thought to encode static body configurations and body parts, whether and how they contribute to the integration of movement information remains an open question.

Unlike general objects, body movements involve complex trajectories of multiple body parts as well as continuous deformation of the overall body shape. Clarifying how motion processing relates to body selectivity in the ventral pathway may therefore provide a key step toward bridging the gap between object perception. Two main mechanisms have been proposed to account for the body movement recognition. The first is the “motion-from-form” hypothesis (Beintema & Lappe, 2002, termed as “shape motion” in the following text), which proposes that body shape and posture are extracted at each moment, and then get integrated into body movements afterwards; and the second is the “optical flow recognition” hypothesis (Giese & Poggio, 2003, termed as “flow motion” in the following text), according to which body movements are recognized from the spatial distribution of the coherent velocity vectors. Unlike shape analysis which can be derived from the static visual features such as body shape or texture, motion pattern analysis treats the motion itself as a defining body feature and relies entirely on the temporal dynamics of the body movements. Accordingly, whereas shape motion can be seen as an extension of classic visual feature hierarchy, flow motion requires a different pathway built on local velocity detectors. These two mechanisms are not necessarily mutually exclusive; therefore, a central question is whether body selective regions encode frame-by-frame static features, transient motion features or both.

There is substantial evidence that regions like EBA and FBA are involved in frame-by-frame body representation. EBA and FBA respond more strongly to videos constructed from frames randomly sampled from different actions than to videos showing intact actions, whereas the motion-selective area, middle temporal cortex (MT), does not show this pattern (Downing et al., 2006). Similarly, Michels et al. (2005) compared classical point-light displays (PLDs; Johansson, 1973) with sequential-position PLDs (Beintema & Lappe, 2002), which eliminate local image motion while preserving form-based structural information. They found robust activations in the FG and pSTS for sequential-position PLD videos, regardless of whether the depicted body structure was moving or static (Michels et al., 2005). Likewise, another fMRI study found that voxel-wise selectivity for PLD body movements in pSTS and EBA was more correlated with selectivity for static body images than with selectivity for coherent dot motion (Peelen et al., 2006). Together, these findings support the view that body-selective regions rely predominantly on shape-based information (Beintema & Lappe, 2002) while motion integration happens elsewhere such as pSTS.

On the other hand, evidence for the recognition of flow motion patterns is scarce, despite a large number of studies conducted on more basic local motion perception in hMT and structure-from-motion perception in dorsal areas (Orban et al., 2006; Orban et al., 1999; Peuskens et al., 2004; Vanduffel et al., 2002). The mechanisms for local motion have been a classic topic in the field of motion perception, termed variously “first-order / low-level motion” (Lu & Sperling, 2001), “surface motion” (Bigelow et al., 2023), or “short-range / local motion” (Braddick, 1974; Hedges et al., 2011). Neurophysiological studies in macaques have revealed distinct brain regions for low-level motion and shape-based motion analysis (Bigelow et al., 2023; Hedges et al., 2011), as well as the encoding of low-level motion features in the dorsal bank of anterior STS body patches (Raman et al., 2023). For higher-level motion patterns, most studies attribute a crucial role to pSTS (Giese & Poggio, 2003; Grossman & Blake, 2002; Jastorff & Orban, 2009; Pitcher, 2025; Pitcher & Ungerleider, 2021; Vangeneugden et al., 2014), while the contributions of the EBA and FBA to flow-motion analysis remain unresolved. Human fMRI studies have revealed substantial anatomical and functional overlap between EBA and MT, including voxels that are selective for body parts as well as for coherent motion (Ferri et al., 2013). Consistent with this finding, joint representation of motion and body shape has been reported in the EBA-homologous body patches of the macaque brain (Bognar et al., 2025). These results suggest that the EBA may also be modulated by flow-motion information. By contrast, the FBA appears to show less prominent motion encoding, both in humans (Robert et al., 2023) and in homologous regions of the macaque brain (Bognar et al., 2025). However, because shape and motion are strongly interdependent, additional empirical controls are needed to rule out potential contributions from structure-from-motion processing.

The present study investigated the contribution of flow motion processing to cortical regions for body processing. We targeted spatiotemporal congruence, an intrinsic property that distinguishes flow motion from shape-motion. In a body movement sequence, each static frame only represents the body features at a single moment, whereas the spatial distribution of velocity vectors are predictive to the motion states of the next moment. Thus, we used synthesized stimuli specifically designed to isolate and modulate the motion vector cues. These stimuli preserved the spatial distribution of transient, pixel-wise coherent motion extracted from naturalistic body movements while eliminating all explicit body shape information. Importantly, these stimuli enabled precise control over both the amount of local motion and its spatiotemporal congruence. Using ultra–high-field fMRI, we examined how motion vector properties influence temporal integration in body-selective areas and how these effects relate to the selectivity for static body shapes, PLD body movements, and coherent dot motion. Furthermore, since it remains questionable whether hard boundaries exist among different categorical clusters (Ritchie et al., 2025), instead of defining the regions of interest (ROIs) separately for each selectivity, we directly quantified shared mechanisms between the flow motion effects and the selectivity through voxelwise correlation within LOTC, FG, and pSTS. We aimed to adjudicate between two competing functional accounts of the role of flow motion in body-movement perception: (1) motion pattern only supplements the recovery of body shape sequences (Beintema & Lappe, 2002; Koerfer & Lappe, 2020), or (2) the spatiotemporal structure of motion pattern itself is sufficient for action recognition (Giese & Poggio, 2003). If flow motion effects reflect shape-sequence analysis, the effects should correlate more strongly with body shape selectivity (Peelen et al., 2006). Alternatively, if flow motion provides independent diagnostic information, the effects should be more closely associated with motion-related selectivity.

## Results

Two experiments were conducted with ultra-high field (7T) fMRI. In the main experiment, we isolated flow motion signals (optical flow, OF) from naturalistic body movements (Supplementary Video 1) using synthesized stimuli that remove explicit body shapes (Figure 1). Each video started with a salt-and-pepper noise frame, and each consecutive frame was generated by displacing the pixels according to the corresponding OF pattern. After adding noise motion to the background, these stimuli preserved the spatial distribution of transient, pixel-wise coherent motion extracted from naturalistic body movements while eliminating all explicit body shape information. The frequency of the presence of body motion was controlled as a factor with three levels, with framewise OFs applied every 2, 4, or 6 frames (15 Hz, 7.5 Hz, and 5 Hz at an overall frame rate of 30 Hz; Supplementary Figure 1). Further, the body-motion optical flows were presented either in their original form (congruent conditions) or after reversing the pixel-velocity directions by 180° (incongruent condition), while preserving their magnitudes and spatial layouts (Supplementary Figure 2). The full experiment had 2 (congruent/incongruent) x 3 (15 Hz/7.5 Hz/5 Hz) = 6 conditions (Supplementary Video 2), presented with an event-related design. The participants were instructed to watch the videos and indicate the walking direction after each video by a button press.

**Figure 1.**
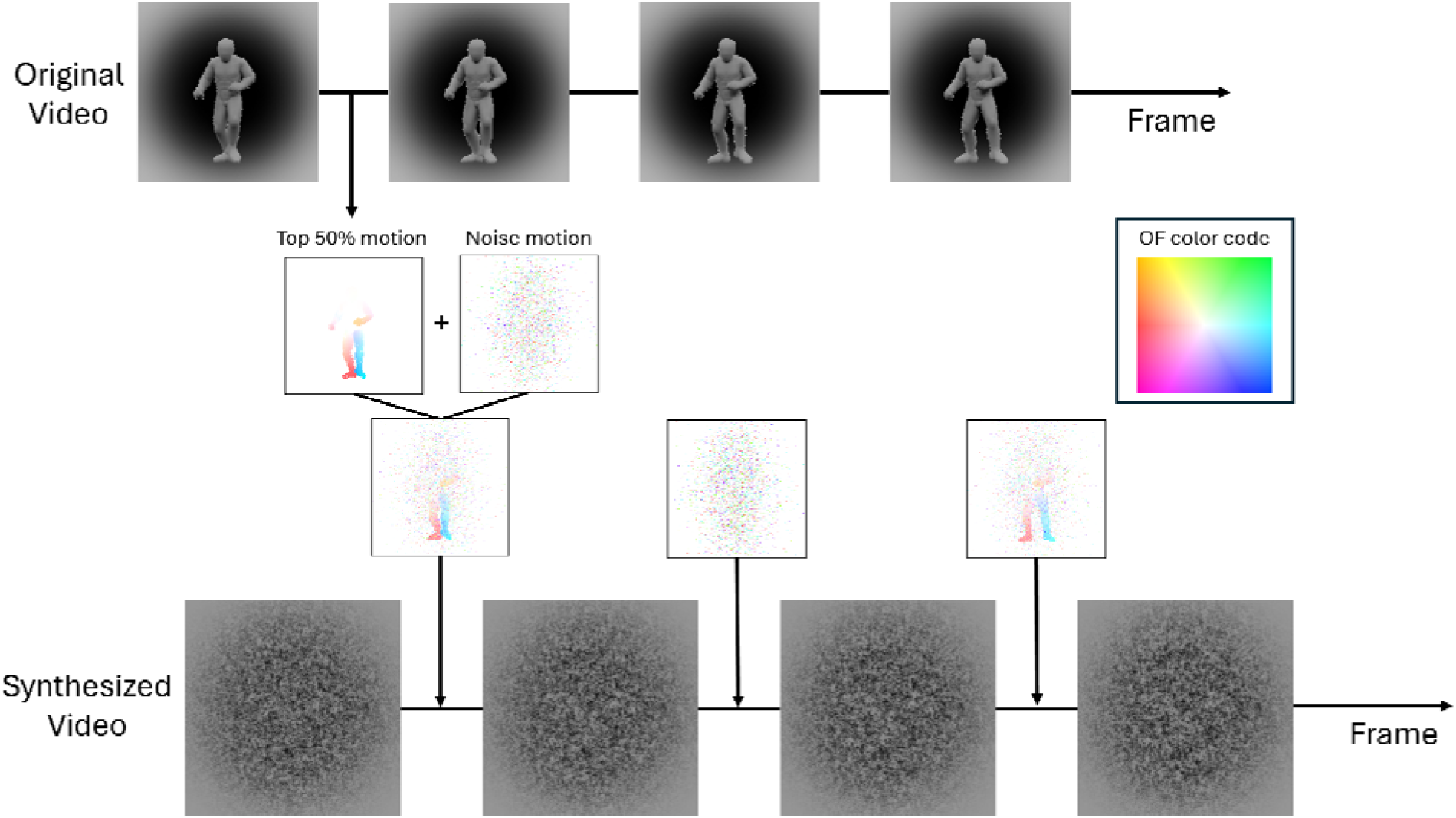
Illustration of OF video synthesizing pipeline. Pixel-wise velocity or optical flow (OF) between each two adjacent frames from the avatar body movement was extracted, filtered to keep the top 50% of motion, and combined with random background motion. A synthesized video started with a salt-and-pepper noise frame. Each subsequent frame was generated by displacing the pixels of the previous frame based on the corresponding OF map. According to different empirical conditions, the applied body motion OF can either be modified or be replaced by random noise motion. Example OF maps are visualized by HSV color code in the figure, with directions presented by the hue and magnitudes presented by the saturation.

A localizer experiment was conducted following the main experiment. Five conditions of videos (PLD body movements, scrambled movements, flickering dots, body images, and scrambled images; Supplementary Video 3) were shown with a block design. Participants were assigned to a passive viewing paradigm with a fixation shape change task to maintain attention.

### Participant exclusion and data quality assessment

The group consisted of thirteen participants. Two participants were excluded from further fMRI analysis due to excessive head motion during scanning. For the rest of eleven participants, we detected suspicious volumes by REFRMS and Global mean amplitude measures. The percentage of marked volumes ranged from 0.66% to 8.86% (mean = 4.10%, SD = 2.44%) for the localizer runs and ranged from 1.12% to 8.14% (mean = 3.60%, SD = 2.25%) for the main experiment runs.

### Regions of interest (ROI) definition and response profiles for the localizer conditions

ROIs were defined for each participant separately. A fixed-effects general linear model (GLM) was performed on the localizer runs to find voxel-wise body motion selectivity. Cortical ROIs were defined by the conjunction contrast analysis of [Body motion > other conditions] and [All conditions > 0]. As a result, three cortical areas were found to be robust across participants: fusiform gyrus (FG), lateral occipitotemporal cortex (LOTC), and pSTS. For FG and LOTC, large clusters were found in all participants for at least one hemisphere and for pSTS in nine out of eleven participants. The spatial coverage of the ROIs is shown in Figure 2. The details of each ROI and each participant are shown in Supplementary Table 3. For bilateral clusters, only the one with higher conjunction t-value was included in the group-level ROI analysis (marked with asterisk in Supplementary Table 3).

**Figure 2.**
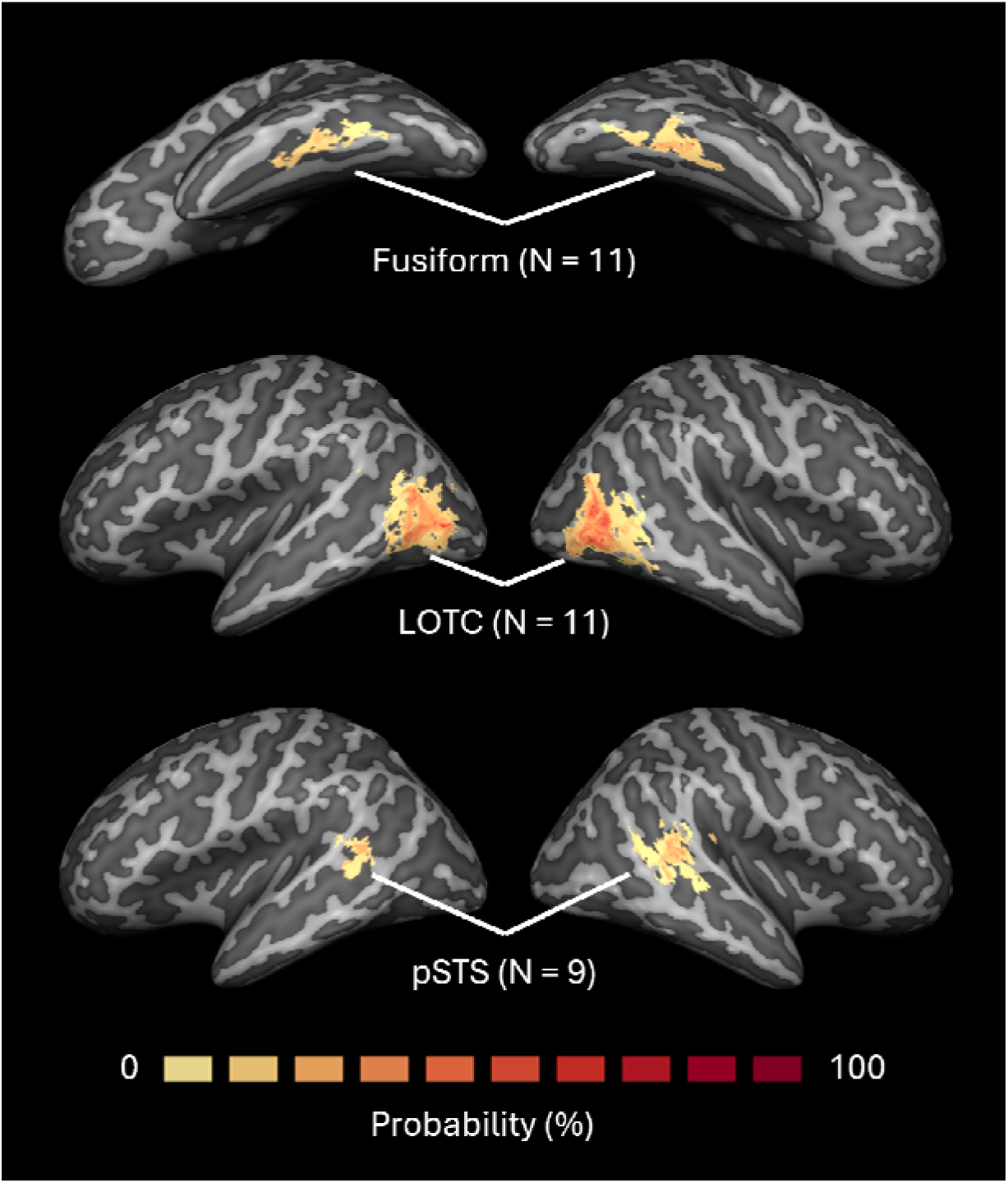
Probability map for subject-level ROIs. The probability maps demonstrate the overlap of significant localizer ROIs across participants. The ROIs were defined at a subject level by the conjunction contrast analysis of [Body motion > other conditions] and [All conditions > 0]. The resulting maps were projected to a normalized brain surface in MNI space and plotted as probability maps across participants. For bilateral clusters, only the one with higher conjunction t-value was included in the group-level ROI analysis (Supplementary Table 3).

A group-level ROI GLM analysis was conducted to investigate the response selectivity for the localizer conditions (Supplementary Figure 3), with contrast analysis tested for body motion selectivity, dot motion selectivity, and static body shape selectivity as shown in Supplementary Table 1. After statistical significance tested by a permutation test with a bootstrapping procedure (Stelzer et al., 2013), body motion selectivity was revealed for all ROIs (FG: N = 11, mean signal change % = 0.786, p < 0.001; LOTC: N = 11, mean signal change % = 0.583, p < 0.001; pSTS: N = 9, mean signal change % = 0.492, p < 0.001). Similarly, dot motion selectivity was also detected in all ROIs (FG: mean signal change % = 0.868, p < 0.001; LOTC: mean signal change % = 1.396, p < 0.001; pSTS: mean signal change % = 0.649, p < 0.001) as well as static body shape selectivity (FG: mean signal change % = 0.715, p < 0.001; LOTC: mean signal change % = 0.571, p < 0.001; pSTS: mean signal change % = 0.347, p < 0.001).

### Behavioral results for the main experiment

The in-scanner behavioral data analysis was conducted with all thirteen participants, including the two excluded from the fMRI analysis (Figure 3). Above-chance accuracy was found for all congruent conditions at three frequency levels (15 Hz: mean = 0.965, p < 0.001; 7.5 Hz: mean = 0.813, p < 0.001; 5 Hz: mean = 0.732, p < 0.001), while significant below-chance accuracy (local motion bias) was found for all incongruent conditions (15 Hz: mean = 0.285, p < 0.001; 7.5 Hz: mean = 0.360, p = 0.009; 5 Hz: mean = 0.363, p < 0.011). Contrast analysis between conditions revealed a significant main effect of the congruence (mean difference = 0.500, p < 0.001) as well as the main effect of the frequency effect (mean difference = 0.078, p = 0.002). The interaction effect was significant (mean difference = 0.311, p < 0.001), showing a significant frequency effect only for the congruent conditions (mean difference = 0.233, p < 0.001) but not for the incongruent conditions (mean difference = -0.059, p = 0.077). All statistical significance was tested by sign-flipping permutation tests at the group-level (3000 repetitions).

**Figure 3.**
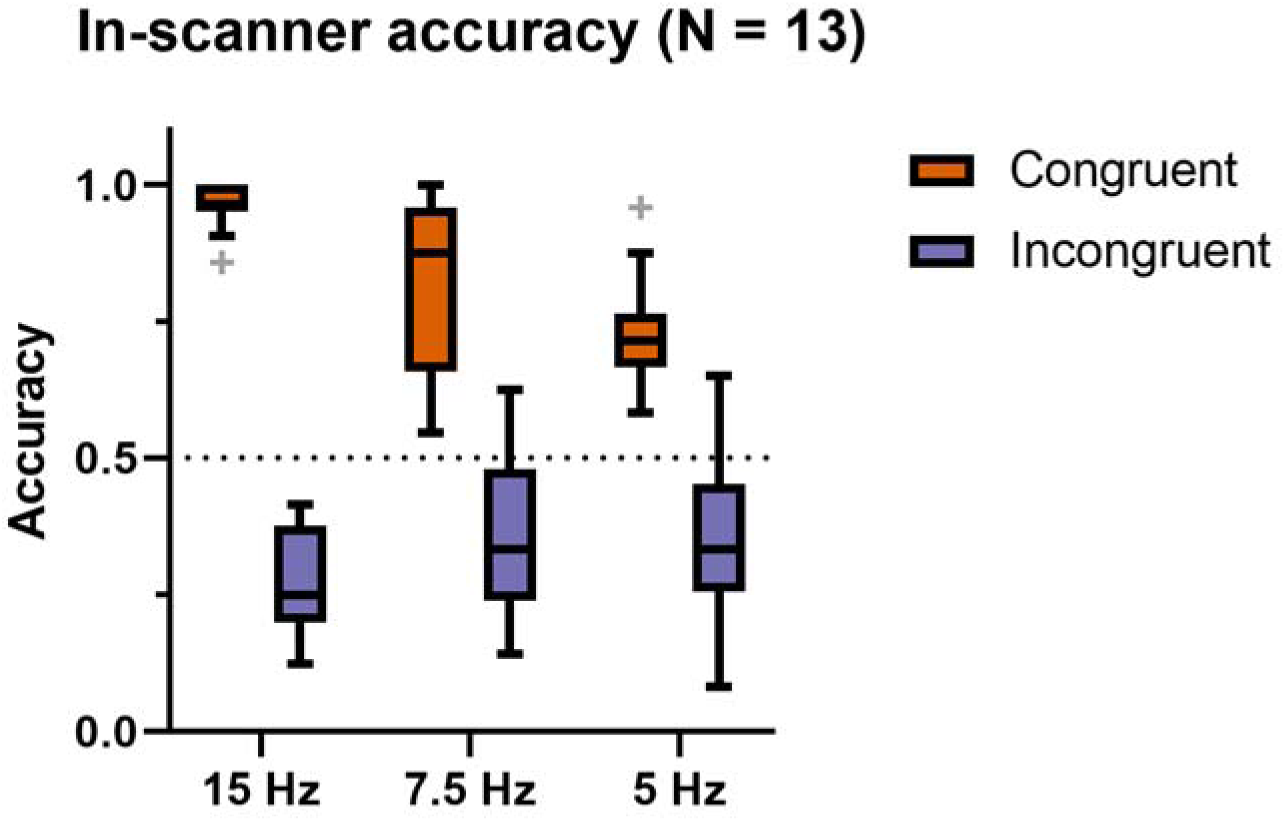
Task performance for OF videos. The boxplots show participants’ accuracies for the in-scanner walking direction task, with whiskers (error bars) represent minimum / maximum values within 1.5 times interquartile range from the lower or upper quartiles and the scatters (+) plotted for individual data beyond the upper and lower bounds.

### ROI response profiles for the main experiment conditions and voxel-wise correlation

A group-level ROI GLM analysis was conducted for the main experiment with contrasts of interest including the congruence effect, the motion frequency effect, and the interaction effect as shown in the Supplementary Table 2. A significant congruence effect was revealed in all three ROIs (Figure 4, Supplementary Figure 4), showing higher responses for congruent conditions than for incongruent conditions (FG: mean signal change % = 0.605, p < 0.001; LOTC: mean signal change % = 0.550, p < 0.001; pSTS: mean signal change % = 0.304, p < 0.001). The frequency effect was also significant for all ROIs, with higher responses for the 15 Hz conditions than for the 5 Hz conditions (FG: mean signal change % = 0.713, p < 0.001; LOTC: mean signal change % = 0.756, p < 0.001; pSTS: mean signal change % = 0.308 p < 0.001). However, the interaction effect was not significant for any of the ROIs (FG: mean signal change % = 0.091, p = 0.290; LOTC: mean signal change % = 0.029, p = 0.544; pSTS: mean signal change % = - 0.073, p = 0.263).

**Figure 4.**
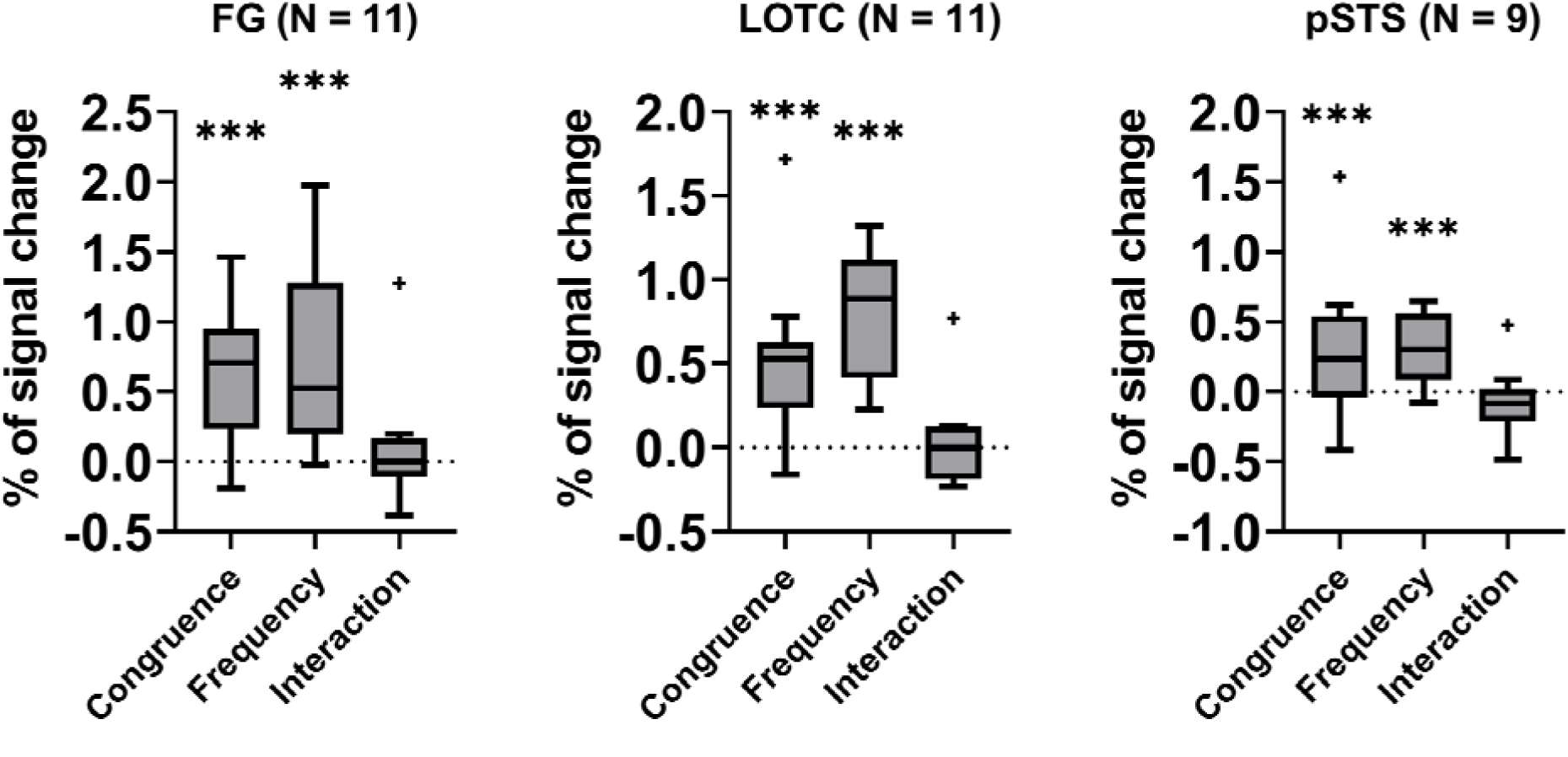
ROI response profiles for flow motion effects. The boxplots show the beta differences resulted from random effect ROI GLM and planned contrasts (Table 2) on the main experiment, with whiskers (error bars) represent minimum / maximum values within 1.5 times the interquartile range from the lower or upper quartiles and the scatters (+) plotted for individual data beyond the upper and lower bounds. For all boxplots, asterisks indicate the significant contrast based on permutation-bootstrapping tests (*: p < 0.05; **: p < 0.01; ***: p < 0.001; two-tailed).

To further clarify the relationship between the main experiment effects and the localizer condition selectivity profiles, we conducted voxel-wise correlation analysis on each ROI (Peelen et al., 2006). Partial correlation was conducted on each ROI’s statistic map between each main experiment effect and the dot motion selectivity, the body shape selectivity, and the body motion selectivity controlled by each other two selectivity maps (Figure 5, Supplementary Table 4). For FG, the congruence effect was positively correlated with the body motion selectivity (mean partial r = 0.268, p < 0.001) and the dot motion selectivity (mean partial r = 0.223, p = 0.004), but was not significantly correlated with the static body shape selectivity (mean partial r = -0.055, p = 0.469). The frequency effect was positively correlated with the body motion selectivity (mean partial r = 0.283, p < 0.001) and the dot motion selectivity (mean partial r = 0.349, p < 0.001) and was weakly negatively correlated with the static body shape selectivity (mean partial r = -0.168, p = 0.026). The interaction effect was not significantly correlated with any of the localizer selectivities (Supplementary Table 4A).

**Figure 5.**
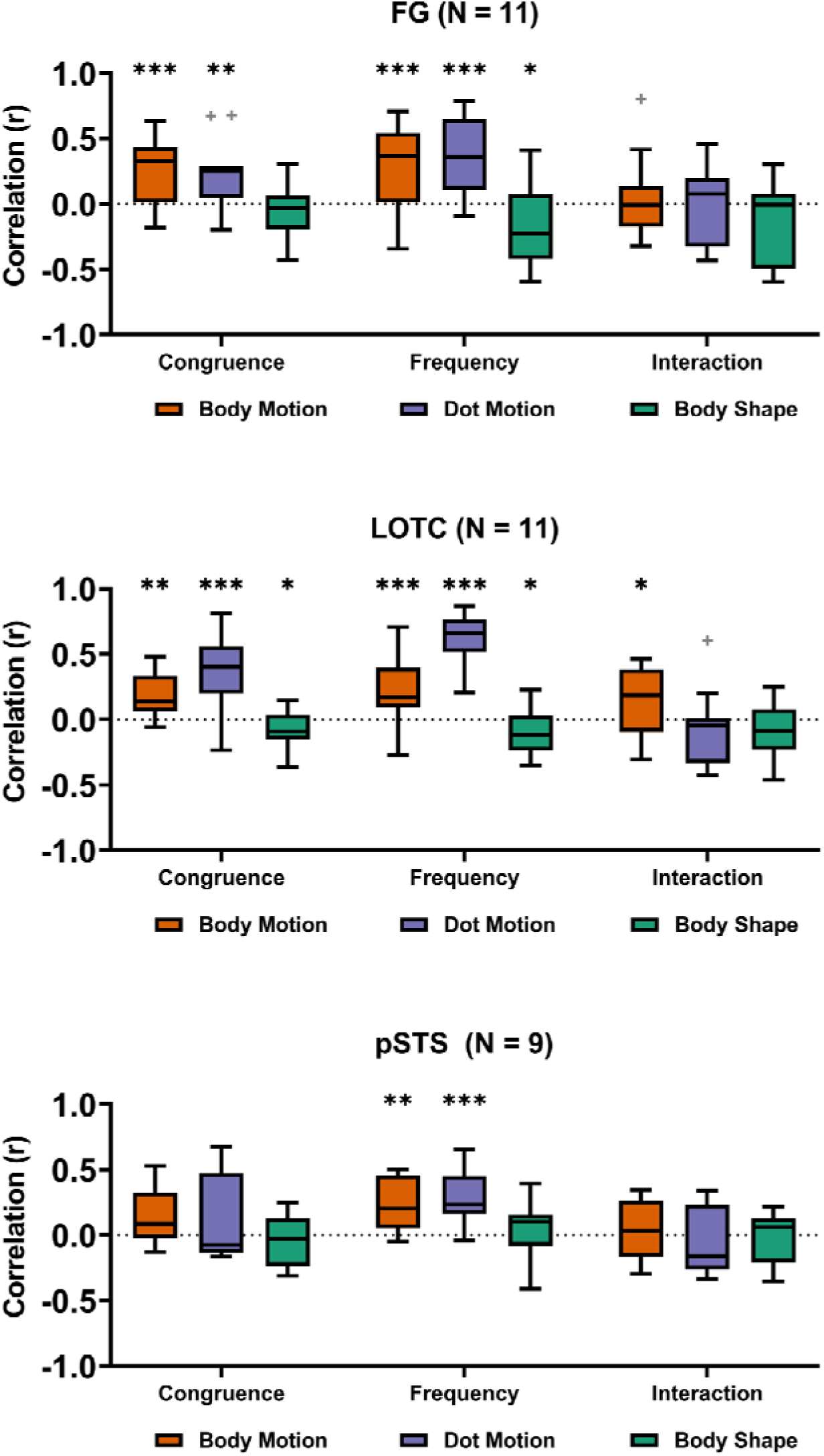
Voxel-wise correlation between flow motion effects and localizer selectivity within each ROI. The boxplots show the partial correlation r-values of voxel-wise correlations between the flow motion effects (congruence, frequency, and interaction) and the localizer-based selectivities (body motion, dot motion, and body shape). Whiskers (error bars) represent minimum / maximum values within 1.5 times the interquartile range from the lower or upper quartiles and the scatters (+) are plotted for individual data beyond the upper and lower bounds. For all boxplots, asterisks indicate significant correlations based on permutation-bootstrapping tests (*: p < 0.05; **: p < 0.01; ***: p < 0.001; two-tailed).

For LOTC, the congruence effect was positively correlated with the body motion selectivity (mean partial r = 0.174, p < 0.005) and the dot motion selectivity (mean partial r = 0.367, p < 0.001), and was marginally negatively correlated with the static body shape selectivity (mean partial r = -0.097, p = 0.043). The frequency effect was positively correlated with the body motion selectivity (mean partial r = 0.204, p < 0.001) and the dot motion selectivity (mean partial r = 0.598, p < 0.001), and was marginally negatively correlated with the static body shape selectivity (mean partial r = -0.096, p = 0.046). The interaction effect was weakly positively correlated with the body motion selectivity (mean partial r = 0.133, p = 0.035), where body motion selectivity was positively correlated with the frequency effect for the congruent conditions (mean partial r = 0.237, p < 0.001) but not significant for the incongruent conditions (mean partial r = 0.088, p = 0.165). The interaction effect was not found to be correlated with the other two localizer selectivities (Supplementary Table 4B).

For pSTS, only the frequency effect was positively correlated with the body motion selectivity (mean partial r = 0.237, p = 0.003) and the dot motion selectivity (mean partial r = 0.253, p < 0.001), but was not significantly correlated with the static body shape selectivity (mean partial r = -0.010, p = 0.882). No significant correlation was detected for the other pairs of effects (Supplementary Table 4C).

## Discussion

We used synthesized pixel-wise optical flow videos to isolate the contribution of shape-free flow motion extracted from naturalistic walking videos. This approach minimized body shape cues and enabled systematic manipulation of temporal congruence across sequences of flow motion cues. Behavioral results showed that the body movement perceived from the synthesized videos was mainly driven by the flow motion instead of the underlying sequence of body structure. At the neural level, multiple cortical areas, including LOTC, FG, and pSTS, were influenced by flow motion properties. In LOTC and FG, voxel responses driven by flow motion were positively correlated with dot motion and body motion selectivity and not with static body shape selectivity. The findings indicate that flow motion serves as a parallel input for body-motion perception in LOTC and FG, offering a new perspective on the cortical pathways underlying body-motion analysis.

### Behavioral Evidence for Local-Motion–Driven Perception

Behavioral results showed that participants could reliably perceive the walking direction from segments of local motion as revealed by significant above chance accuracy for the congruent condition. In contrast, when the flow motion velocity was flipped in incongruent conditions, accuracy dropped below chance, even though the sequence of implied body postures remained unchanged. These findings confirmed that the recognition of movement direction in the synthesized videos was not driven by the trajectory of motion-induced body shapes but instead, was derived from the flow motion patterns.

We further investigated how the spatiotemporal dependence of flow motion segments may affect the integration of global body movements. A familiar assumption is that the perception of global motion requires the accumulation of segmented motion over time (Blake & Shiffrar, 2007; Oguz et al., 2024)). If so, task performance should be dependent on the total amount of motion presented and the accumulation of motion signals would fail in the incongruent setting, as the motion information between consecutive segments is inconsistent. Thus, we compared the task performance between videos with the lowest (5 Hz) and the highest (15 Hz) amount of body motion. Increasing the amount of motion improved task accuracy for the congruent conditions but not for the incongruent conditions, with a significant interaction effect observed between congruence and motion frequency. Together with the below-chance accuracy, the results suggest that in incongruent conditions, participants’ decisions were biased by the individual motion segments and not based on integrated body motion.

### Shared mechanism between flow motion and body movement processing in FG and LOTC

For FG, LOTC, and pSTS, we found significant higher activations for congruent motion conditions than for incongruent ones. Since the congruent videos had the same underlying body shape sequences and total motion energy as the incongruent ones, the congruence effect observed in the ROIs reflects low-level modulations by the flow motion signal. To further investigate the contribution from the shape-driven motion processing, we conducted partial correlation analysis between the flow motion effects and the localizer selectivity. Various flow motion effects were positively correlated with body-motion and dot-motion selectivity, whereas they showed nonsignificant or marginal negative correlations with body shape selectivity. The results contrast with Peelen et al. (2006), where the conventional PLD body motion was strongly correlated with the body shape selectivity instead of motion selectivity. Thus, the current results separated the flow motion effects from the motion-from-shape mechanism (Beintema & Lappe, 2002). Instead, the results are in line with the computational framework of the optical flow motion pathway as proposed by Giese and Poggio (2003), which predicted the interference of motion integration due to the inconsistency between segments of the motion signal.

Our synthesized stimuli with minimal shape information provided the opportunity to revisit the roles of conventional body selective areas in body motion processing. We first found significant main effects of congruence and motion frequency in FG. Both effects significantly correlated with body motion and dot motion selectivity at the voxel level, while no interaction related effects or correlations were found. Thus, the two flow motion effects (congruence and frequency) and the two localizer selectivity (body motion and dot motion) results may be driven by similar factors. One candidate factor may be the general articulated structures implied by the motion pattern (Taylor & Downing, 2011; Taylor et al., 2007). In the main experiment, inducing motion inconsistency can introduce ambiguity of the underlying structure through the interrupted global motion signals, while reducing the motion frequency directly decreases the amount of the shape-from-motion cues. Meanwhile in the localizer experiment, the structural cues decreased from the preferred one as in the body motion condition, to the random non-preferred structure as in the scrambled dot motion condition, and finally to the absence of structure as in the flickering dot condition. However, the configuration cues in our stimuli were derived from motion rather than from static body shapes, challenging the view that FG analyzes only individual frames, i.e. static body shape, rather than the overall temporal dynamics (Downing et al., 2006). Furthermore, the absence of a correlation with body shape selectivity indicates that the motion inputs served a role beyond merely cueing body shape segmentation. Thus, the current finding suggested a significant contribution of FG to the local coherent motion pattern analysis.

Within LOTC, dot motion selectivity was correlated with both the congruence effect and the motion frequency effect. Because LOTC region encompasses multiple motion selective areas including hMT, the dot motion correlation observed here may reflect differences in global and local motion energy manipulated by the motion consistency and temporal frequency. More revealing, however, were the patterns related to body-motion selectivity. Voxelwise body motion selectivity in LOTC was strongly correlated to the frequency effect only under congruent conditions but not under the incongruent ones. Together with LOTC’s preference for congruent motion, this pattern suggests that body motion selectivity of the LOTC area reflects the spatiotemporal integration of the global motion pattern, a process that is disrupted when flow motion segments are temporally incongruent. Importantly, rather than subdividing LOTC into hMT+ and EBA, we leveraged voxelwise selectivity to capture the functional heterogeneity across the LOTC region. These results may help explaining the prior reports of limited motion contributions in the static body defined EBA (Grossman & Blake, 2002; Peelen et al., 2006): we observed slightly negative correlations between flow motion effects and static body shape selectivity, indicating that the voxels most responsive to static shape feature were less modulated by the flow motion cues. Together with the findings in FG, these results argue against a strict hierarchical account, in which motion signals are sole extracted from body shape sequence. Instead, they support a model in which low-level motions and body shapes are encoded as partially separable dimensions within FG and LOTC.

Surprisingly, although pSTS is well known to be involved in body motion processing (Grossman et al., 2005; Grossman & Blake, 2002; Jastorff & Orban, 2009; Vangeneugden et al., 2014), it showed no correlation between the flow motion effects and the body motion selectivity. Despite the pSTS preference toward the congruent conditions, the absence of correlation between the congruence effect and the body motion selectivity suggested that the PLD body motion selectivity may be driven by a different mechanism. Nevertheless, the results are well predicted by the study of Michels et al. (2005), in which the sequential position PLD was used to remove the low-level motion. In addition to the classic body motion selectivity, comparably high activation was found in pSTS for sequential position PLDs with static body postures (Michels et al., 2005). Thus, instead of motion itself, the pSTS may be involved in estimating and updating the body shape across time. These results underscored the importance to separate the motion features when studying body movement perception. While the PLD format is efficient for the laboratory control of visual features, it oversimplifies the naturalistic body movements by sacrificing the feature-rich motion field. This simplification may introduce redundant body structure computations to articulate the isolated joint points before extracting the global motion pattern (Duarte et al., 2022; Hirai & Senju, 2020). Furthermore, due to the loss of motion features, such modification may misrepresent the cortical basis for naturalistic body movement recognition especially by overestimating the body shape contribution.

### Reconsidering hierarchical models of body perception

Here, by taking the flow motion signal as the primary visual input, we observed the sensitivity in LOTC to the global body movement. In line with the neuroimaging results, our behavioral results revealed stronger bias toward the local motion instead of the shape-from-motion sequence, suggesting that the computation within LOTC may serve as the major source for higher-level decision making in this task. Moreover, our data suggested that FG may analyze transient flow motion patterns for an intermediate stage of motion processing, analogous to the computations of “OF pattern neurons” proposed by Giese and Poggio (2003). Thus, one remaining question is how this motion pathway may support the higher-level recognition for the body movements. In addition to the parallel processing view (Giese & Poggio, 2003), an alternative hypothesis is that body movement perception is achieved in a hierarchical manner, with FG and LOTC first detecting body forms and pSTS integrating these signals into higher-level perceptions (Grossman & Blake, 2002; Jastorff & Orban, 2009; Landsiedel et al., 2022). However, the current study indicates that the processing of isolated flow motion cues can be already detected in FG and LOTC. In line with this result, other studies have reported that multiple categorical dimensions embedded in the low-level motion field can be decoded from temporal occipital areas (Robert et al., 2023; Schmid et al., 2021). These results are compatible with the “third pathway” (Pitcher & Ungerleider, 2021), which proposed a motion-driven pathway through hMT+ and STS. Future studies may focus on whether those motion-based analyses will be merged into pSTS, or instead, will be directly read out through other parallel cortical pathways (Sokolov et al., 2018).

Another key question is whether the integration between shape and motion occurs within LOTC. Previous studies have shown that LOTC represents multiple shape-based body features including joint positions (Marrazzo et al., 2023), whole-body postures (Poyo Solanas et al., 2020), and biomechanical dynamics (Marrazzo et al., 2025). While it has been proposed that the body movements are encoded in LOTC as “snapshots” or discrete frames (Downing et al., 2006), it remains unclear if these frame-wise representations can be modulated by the motion cues. Importantly, LOTC responses have been found sensitive to the temporal context. Temporally scrambled videos (Smekal et al., 2025) and videos with random body postures (Downing et al., 2006) elicit stronger activations than coherent, continuous body movement in LOTC. This pattern is compatible with the suppression effects observed in early visual cortex for shape-based apparent motion (Alink et al., 2010; Hidaka et al., 2011; Shen et al., 2020), which predicts stronger activation for violations to temporal contexts. However, such mechanism is incompatible with the flow motion effects found in the current study, where the inconsistency across time resulted in decreased activities. This dissociation suggests that LOTC computations flow motion differently from shape-based motion. Conceptually, shape-based motion analysis requires integrating multiple position changes across frames to infer global movements, the flow motion reflects the motion state at each individual position (Kwon et al., 2015). Furthermore, the flow motion states may also affect the efficiency when tracking the trajectories of multiple body parts (Clair et al., 2010; Kwon et al., 2015) and estimate the body part position (Maus et al., 2013; Nishida & Johnston, 1999; Ramachandran & Anstis, 1990; Whitney, 2002). Interactions between static features and implied motion have been reported in LOTC (Kourtzi & Kanwisher, 2000; Lorteije et al., 2006; Pavan et al., 2011). An important extension of the present work would be to test whether flow motion states modulate LOTC encoding of static body postural features (e.g. joint position), for example by overlaying OF motion onto static body shapes.

Finally, the current results raise the question about the subcortical contributions for the body movement processing. Representations for complex motion patterns have been found in multiple subcortical structures including superior colliculus (SC; Lu et al., 2024; Schneider & Kastner, 2005; Wolf et al., 2015) and pulvinar (Cotton & Smith, 2007; Villeneuve et al., 2005), which have anatomical connectivity to hMT+ (Arcaro et al., 2015; Lanyon et al., 2009). Studies with fMRI and tractography have proposed the pathway between LGN and hMT+ that bypasses V1 and serves as the entrance to cortical process for motion (Ajina & Bridge, 2018; Ajina et al., 2015; Gaglianese et al., 2012). These observations raise the possibility that motion signals may access body-sensitive cortex through pathways that are at least partly distinct from those carrying detailed shape information. Clarifying how subcortical and cortical motion pathways interact—and whether they converge or remain segregated before reaching body-selective regions—will be essential for understanding the neural architecture supporting biological motion perception and for establishing functional homologies between human and non-human primate systems.

## Methods

### Participants

Thirteen healthy participants (age = 21.31 ± 2.66 years; 4 males, all right-handed) participated in the experiment. All participants had normal or corrected-to-normal vision and no medical history of any psychiatric or neurological disorders. During the recruitment, the original body movement videos (Supplementary Video 1) were provided to the potential participants. Only participants who could clearly differentiate the walking directions were finally included into the experiment. All participants provided informed written consent before the start of the experiment and received a monetary reward (vouchers) or course credits for their participation. The experiment was approved by the Ethical Committee at Maastricht University and was performed in accordance with the Declaration of Helsinki.

### Main experiment

#### Stimuli

##### Original animation and optical flow extraction

The original video was downloaded and edited from Adobe’s Mixamo animation library (https://www.mixamo.com/). The video depicted a human actor moving to the right while keeping the upper body facing forward (“Jog Strafe Right”). We chose this specific strafe animation because it can also be played backward to present a leftward moving pattern. Thus, the forward- and backward-played videos contain the same set of body postures, while the moving direction can only be inferred from the motion features. The full video presented three walking cycles in 3.6 seconds at a frame rate of 30 Hz.

The original animation was further edited in Blender (https://www.blender.org/), including removing the global translation, reducing keyframes, and enlarging the body area. A rendered video is shown in Supplementary Video 1 with a frame rate of 30 fps. After the editing, the framewise optical flow (OF) patterns were extracted during rendering using a third-party plugin in Blender (Cartucho et al., 2021). For each frame, the motion information was removed from 50% of the pixels with the lowest speed. This ensured that only the most salient local motion was preserved in the OF patterns.

To reduce the segmentation of body shape from motion against the static background, random background motion was added to each OF pattern. For each individual OF pattern, the original body area was first binarized and then filtered with a 2D low-pass filter with a cutoff spatial frequency of 2 cycles/image. The filtered image then served as a probability map for adding random motion, resulting in a noise distribution with higher density closer to the body area and lower density in the periphery of the visual field. The body OF was then overlaid on top of the noise motion and used as a reference to synthesize the body motion videos.

##### Video synthesizing procedure

The synthesized videos were generated by continuously applying the OF patterns frame by frame (Figure 1). Each video started from a salt-and-pepper noise image, and each consecutive frame was generated by applying the corresponding OF pattern as the pixel-wise displacement map to the current frame. Importantly, the low-level motion was constrained to the static body area in each frame, so that the generated motion reflected only local pixel movements rather than actual body-part displacements. The high spatial frequency components (> 55 cycles / image) were removed from each generated frame to reduce the sharp boundary of each pixel.

We created different experimental conditions by modifying the reference OF patterns. The first configuration controlled the amount of presented body motion (Supplementary Figure 1). Instead of applying the full sequence of the walking OFs, framewise OFs were applied every 2, 4, or 6 frames, interleaved with the noise motion OF described above. Thus, instead of a continuous body motion flow, each video consisted of a discrete sequence of transient local motion segments. At an overall frame rate of 30 Hz, the temporal frequency of body motion presentation became 15 Hz, 7.5 Hz, and 5 Hz.

Secondly, the body-motion optical flows were presented either in their original form or after reversing the pixel-velocity directions by 180°, while preserving their magnitudes and spatial layouts (Supplementary Figure 2). Unlike the original flows, the reversed flows disrupted the spatiotemporal continuity of motion across successive local motion segments. We hypothesized that such incongruence may further impair the integration of a local motion sequence into global motion perception. Moreover, because the spatial distribution of moving pixels remained the same between congruent and incongruent versions, the sequence of motion-induced shape was well-controlled. This configuration, therefore allowed us to isolate local motion modulation while controlling for residual shape-induced motion.

Finally, the synthesized frames were combined in either forward or backward-played order. Since the original animation depicted a rightward walking pattern, the backward-played version showed a leftward direction instead. Except for this walking direction difference, the backward videos with congruent OF motion contained the same set of motion segments as the forward videos with incongruent OF motion. Thus, by pooling forward and backward trials, the effects of motion direction were balanced between congruent and incongruent OF conditions.

All synthesized videos were resized to 1200 x 1200 pixels underneath a circle aperture spanning 10 degrees in visual angle. The overall body area was 7 x 5 degrees in visual angle. Examples of synthesized videos are shown in Supplementary Video 2.

#### Experimental design

The experiment had 2 (congruent/incongruent) x 3 (15 Hz/7.5 Hz/5 Hz) = 6 conditions, presented with an event-related design. Each trial started with a video of one condition playing for 3.6 s, either forward-played or backward-played, followed by a jittered intertrial interval of 10.7 s, 12 s, or 13.3 s. The participants were instructed to watch the videos and indicate the walking direction after each video by a button press. A white “+” fixation (0.5 degree visual angle) was overlaid on the center of the display throughout the experiment. It turned to green after each video as a signal for the participant to respond and turned back to white after the button press was received. No feedback was given regarding the response correctness during the experiment.

For each main experimental run, each condition was repeated eight times (four forward-played and four backward-played) and presented in a pseudorandom order, resulting in 48 trials spanning thirteen minutes. Each participant finished three main experimental runs, resulting in 144 trials (24 repetitions per condition) in total.

### Localizer experiment

#### Stimuli

PLD body motion videos were generated from five different whole-body movements, adapted from Mixamo (https://www.mixamo.com/) animations including ball throwing, dancing, and kicking. All selected animations were cut to a length of 1 second, centered to the field of view, and duplicated with a horizontally mirrored version, resulting in ten unique videos in total. For each video, trajectories of joint markers (head, neck, shoulders, elbows, wrists, hands, hop, groins, knees, ankles, and feet) were extracted and formatted into a PLD animation at a frame rate of 30 Hz.

Based on the joint dynamics of PLD body motion, scrambled motion and flickering dot videos were created. The scrambled motion videos maintained the trajectory of each joint point while randomizing the onset horizontal position of the joint points. Thus, in the scrambled motion condition, the overall motion energy remained the same while the body configuration was disrupted. In the flickering dot condition, in addition to the onset randomization, the motion was further removed by randomly displacing half of the joint points in a random direction and keeping the rest static in each consecutive frame.

Two more static image conditions were created to test body shape selectivity. The first and middle frames of each body motion video were drawn as stick figures, resulting in twenty different images displaying a static body posture. As a control, scrambled images were generated by inverting the stick figures, randomizing the onset horizontal position the joint points and randomizing the joint point connections, resulting in twenty corresponding abstract figures with no body configuration.

All videos were resized to 1200 x 1200 pixel spanning 10 degrees in visual angle. The overall stimulus area was 7 x 5 degrees in visual angle. Examples of localizer stimuli are shown in Supplementary Video 3.

#### Experimental design

The functional localizer used a blocked design with five conditions of videos / images. Within each video condition block, five videos were presented for 1 second each and separated by a fixed 200-ms inter-trial interval. And for each image condition block, ten images were presented for 500 ms, interleaved by a fixed 100-ms inter-trial interval. The total length of each block was 6 seconds for all conditions. The order of block conditions was randomized for each participant, and each condition was repeated nine times within each run. Two consecutive blocks were interleaved by a jittered inter-block interval of 9.6, 10.9, or 12.2 seconds. A white “+” fixation was overlaid at the center of the display throughout the run. Each run contained five catch blocks (one catch per condition) where, in one of the trials, the fixation point changed its shape from a “+” to an “x”. Participants were instructed to press a button when they detected a change of the fixation shape. Each localizer run lasted for 13.5 minutes, and each participant finished two runs (18 repetitions per condition) during the scanning.

### Stimulus presentation

All experiments were programmed using Psychtoolbox (https://www.psychtoolbox.net), implemented in Matlab 2018b (https://www.mathworks.com). Stimuli were projected onto a screen positioned at the end of the scanner bore with a Panasonic PT-EZ57OEL projector (screen size = 30 x 18 cm, resolution = 1920 * 1200 pixels). Participants viewed the stimuli through a mirror attached to the head coil (screen-to-eye distance = 99 cm, visual angle = 17.23 x 10.38 degrees).

### MRI acquisition

All images were acquired with a Siemens 7T MAGNETOM scanner with a 1-transmitter/32-receiver head coil (Nova Medical) at the Maastricht Brain Imaging Centre (MBIC) of Maastricht University, the Netherlands. Functional images were collected using the T2*-weighted multi-band accelerated EPI 2D BOLD sequence (TR/TE = 1300/20 ms; multiband acceleration factor = 3; flip angle = 59°; in-plane isotropic resolution = 1.6 mm, number of slices per volume = 66; matrix size = 128 x 128; volume number = 594 for the main experiment and 603 for the localizer). T1-weighted anatomical images were obtained using the 3D-MP2RAGE sequence (TR/TE = 5000/2.47 ms; Inverse time TI1/TI2 = 900/2750 ms; flip angle FA1/FA2 = 5/3°; in-plane isotropic resolution = 0.7 mm; matrix size = 320 x 320; slice number = 240).

### Data analysis

#### fMRI preprocessing

For anatomical images, preprocessing was conducted with a third-party SPM pipeline (PreSurfer; Kashyap et al., 2021) under Matlab 2018b. Steps included MP2RAGE denoising, biasfield correction, and brain stripping. The resolution was then downsampled to 0.8 mm for better alignment to the 1.6 mm resolution of functional images. For functional images, the preprocessing steps were conducted by Brainvoyager 22.2 (https://www.brainvoyager.com/), including EPI distortion correction (Breman et al., 2020), slice scan time correction, 3D head motion correction, and high-pass temporal filtering (GLM with a Fourier basis set of 6 cycles / run, including linear trend). Trilinear/sinc interpolation was used in the motion correction step, and sinc interpolation was used in all other steps.

Quality assessment was conducted on each preprocessed functional run using two additional measures: the residual mean square error to reference frame (REFRMS) and the global mean amplitude change over time. The REFRMS algorithm was adapted from the “fsl_motion_outliers” function implemented in FSL (https://web.mit.edu/fsl_v5.0.10/fsl/doc/wiki/FSLMotionOutliers.html), without calculating the REFRMS difference between consecutive frames. The measure thus quantifies the normalized intensity difference between each volume and the first volume, accounting for the residual errors after head motion correction. Similarly, global mean amplitude was calculated for each volume, accounting for the overall signal changes due to head motion or magnetic field inhomogeneity across time. The two measures were stored and served as indices for volume censoring in the later stage data analysis.

The first volume of the first functional run was chosen as a reference image for the following coregistration procedure. Displacement fields were estimated for the first volumes of all other functional runs with SyN deformation implemented in ANTs toolbox (Avants et al., 2009), and further corrected with Lanczos-windowed-sinc interpolation. The preprocessed anatomical image was also warped to the reference functional image to account for the residual EPI distortion, resulting in a warped T1-weighted anatomical image. The aligned functional and anatomical images were then normalized to MNI space based on the SyN deformation estimated between the warped anatomical image and the MNI reference image. No spatial smoothing was applied in any of the steps to preserve the fine-grained voxel responses.

#### Noise and suspicious volume estimation

Physiological noise was estimated for each functional run using the CompCor algorithm (Behzadi et al., 2007). Each participant’s normalized anatomical image was segmented with a deep learning based pipeline (FastSurfer; Henschel et al., 2020) under FreeSurfer (Fischl, 2012). White matter and CSF masks were generated and eroded for 2 mm to extract non-gray matter signals. Five components were extracted for each mask and for each run with the aCompCor approach (Behzadi et al., 2007), then orthogonalized against the experiment design, resulting in ten independent confound time-courses. In addition, for each run, suspicious volumes were marked based on the REFRMS and global amplitude measures using box plot outlier limits for each measure.

#### Subject-level regions of interest (ROI) definition

ROIs were defined for each participant separately. A fixed-effects general linear model (GLM) was performed on the localizer runs to find voxel-wise body motion selectivity. In the design matrix, each condition predictor was modeled as a boxcar function with the same duration of the block and convolved with the canonical hemodynamic response function (HRF). CompCor and motion confounds were added as nuisance regressors. Cortical ROIs were defined by the conjunction contrast analysis of [Body motion > other conditions] and [All conditions > 0]. No volume censoring was applied at this stage. The resulting statistical map was corrected using a cluster-threshold statistical procedure based on Monte-Carlo simulation (initial p < 0.001, alpha level = 0.05, iterations = 3000). The clusters were grouped based on each participant’s anatomy. For bilateral clusters, only the one with higher conjunction t-value was included in the group-level ROI analysis.

#### Localizer ROI analysis

Group-level analysis was conducted on the localizer runs to further investigate each ROI’s response profile. For each participant and each ROI, an ROI time-course was extracted by averaging the voxel time-courses and was normalized to percentage of change. A fixed-effects GLM at subject-level was conducted on the extracted ROI time-courses with the localizer conditions as predictors and the CompCor/head motion as confounds. Volume censoring was applied before GLM by removing each run’s outlier volumes from both the time-course and from the design matrix. Contrast analysis was conducted on each participant’s estimated betas for body motion selectivity, dot motion selectivity, and static body shape selectivity as shown in Supplementary Table 1. The beta differences were then averaged across participants, resulting in a group mean beta difference for each contrast.

Group-level statistical significance was tested by a permutation test with a bootstrapping procedure (Stelzer et al., 2013). At the subject-level, the condition labels of blocks were shuffled within each run, after which the fixed-effects GLM and the contrast analysis were recalculated. The procedure was repeated for 2000 times per participant, resulting in a subject-level null-distribution of the contrast differences. Next, at group-level, bootstrapping was conducted on the participants by randomly drawing one sample with replacement from each participant’s null-distribution and recalculating the group mean. This procedure was repeated 10000 times, generating a group-level null-distribution of the contrast beta difference. The p-values of the contrasts were calculated as the percentage of the absolute values of bootstrapped mean that are larger than the absolute value of experimental mean (two-tailed test).

#### Main experiment behavioral data analysis

The group-level statistics were first conducted in a non-parametric way to test the accuracy of each condition against the chance accuracy of 0.5. The accuracy of each condition was averaged across participants after subtracting the chance accuracy. A permutation null-distribution was generated by recalculating the group mean for 3000 times after randomly flipping the sign of each participant’s accuracy difference from chance. The p-values of the contrasts were calculated as the percentage of the absolute values of bootstrapped mean that are larger than the absolute value of experimental mean (two-tailed test).

The accuracy differences between conditions were then tested with three planned contrasts (Supplementary Table 2), including the main effect of congruence, the main effect of motion frequency, and the interaction between the two factors. The accuracy difference was calculated for each contrast and averaged across participants. Permutation test was conducted for each contrast separately following the sign-flipping approach, resulting in a null-distribution with 3000 samples each for the two-tailed significance test.

#### Main experiment ROI analysis

The group-level ROI analysis for the main experiment followed the same pipeline as in the localizer ROI analysis. The contrasts of interest for the main experiment included the congruence effect, the motion frequency effect, and the interaction effect as shown in the Supplementary Table 2. Permutation and bootstrapping steps were conducted to test the statistical significance, with 2000 samples for each subject-level null-distribution and 10000 samples for the group-level null-distribution.

#### Voxel-wise correlation

To further clarify the relationship between the main experiment effects and the localizer condition selectivity profiles, we conducted voxel-wise correlation analysis on each ROI (Peelen et al., 2006).For each participant and each ROI, a voxel-wise fixed-effects GLM was conducted on localizer and main experiment runs, followed by the corresponding contrast analysis as shown in (Supplementary Table 1 & 2). The resulting beta differences for each contrast were z-scored across voxels within each ROI. Partial correlation was conducted on each ROI’s statistic map between each main experiment effect and the dot motion selectivity, the body shape selectivity, and the body motion selectivity controlled by each other two selectivity maps. The subject-level partial correlation r-values were averaged across participants for each corresponding ROI and entered into a group-level statistical test.

The group-level statistics followed the permutation and bootstrapping approach as mentioned above. First, the GLM and ROI statistic maps were recalculated for the main experiment runs after shuffling the trial labels within each run. The partial correlation was recalculated for each pair of the maps, resulting in subject-level null-distributions of r-values, with 2000 samples generated for each pair of maps. Next, group means of the r-values were calculated after bootstrapping the subject-level null-distribution for 10000 times, resulting in the group-level null-distributions. A two-tailed test was conducted between each experimental r-value and the corresponding group null-distribution.

