## Supplementary Figures & Tables for "Optical flow analysis dissociates cortical selectivity for body movements and shapes"

Local Motion as a dissociable dimension for Human Body Movement Selectivity in High-Level Visual Cortex

Supplementary Figures

### Supp. Figure 1. Illustration of applied OF sequence in different frequency settings.

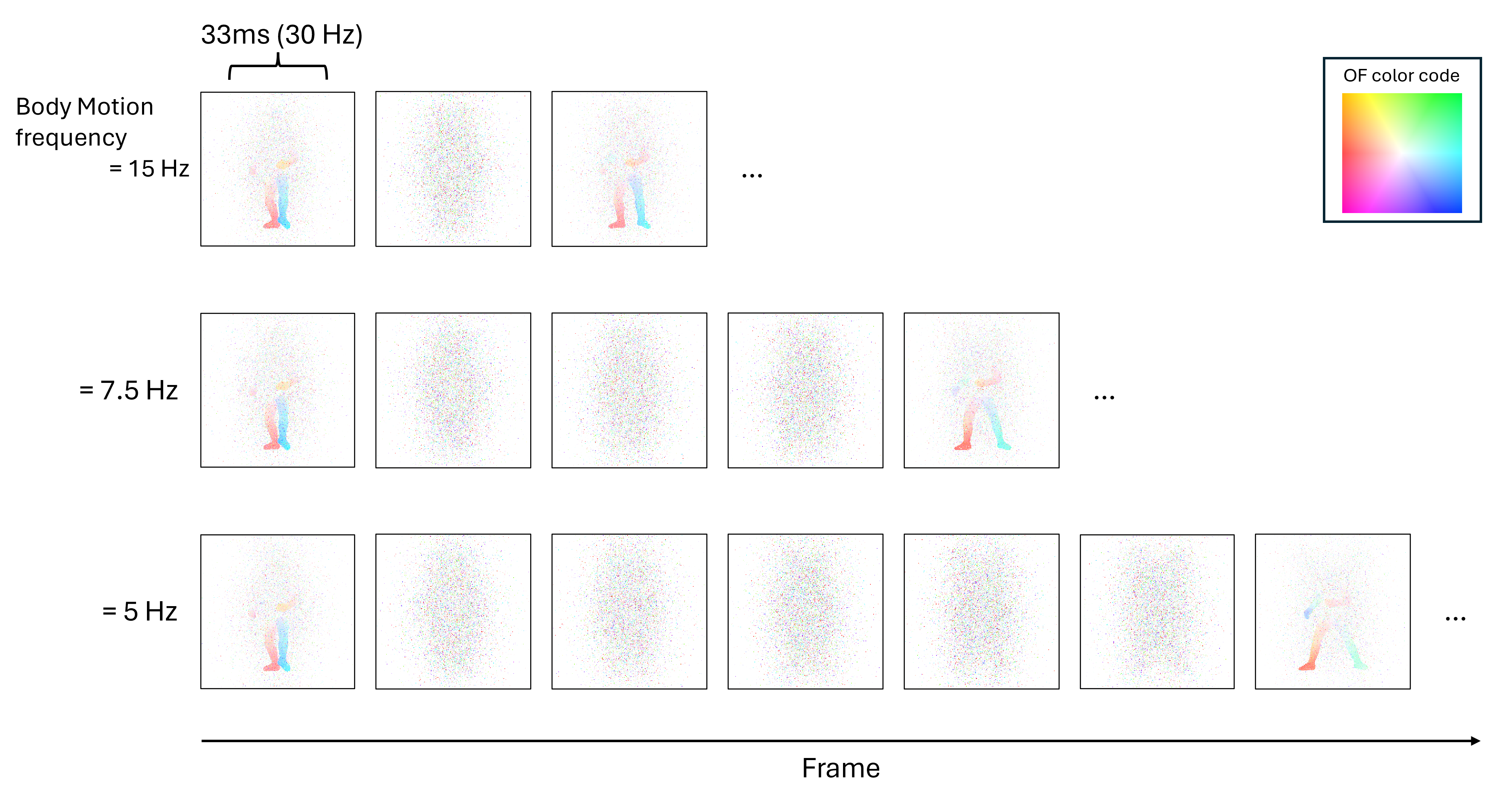

The pixel-wise OF motion was applied to generate each subsequent frame, creating a transient motion (33 ms) between each two frames at a frame rate of 30 Hz. The OF motions contained either body motion patterns or only noise motion. The body motion OFs were applied every two, four, or six frames, interleaved by noise motions. At a frame rate of 30 Hz, the presence of body motion became 15 Hz, 7.5Hz, and 5 Hz accordingly.

Supp. Figure 2. Illustration of applied OF sequence in different congruence settings.

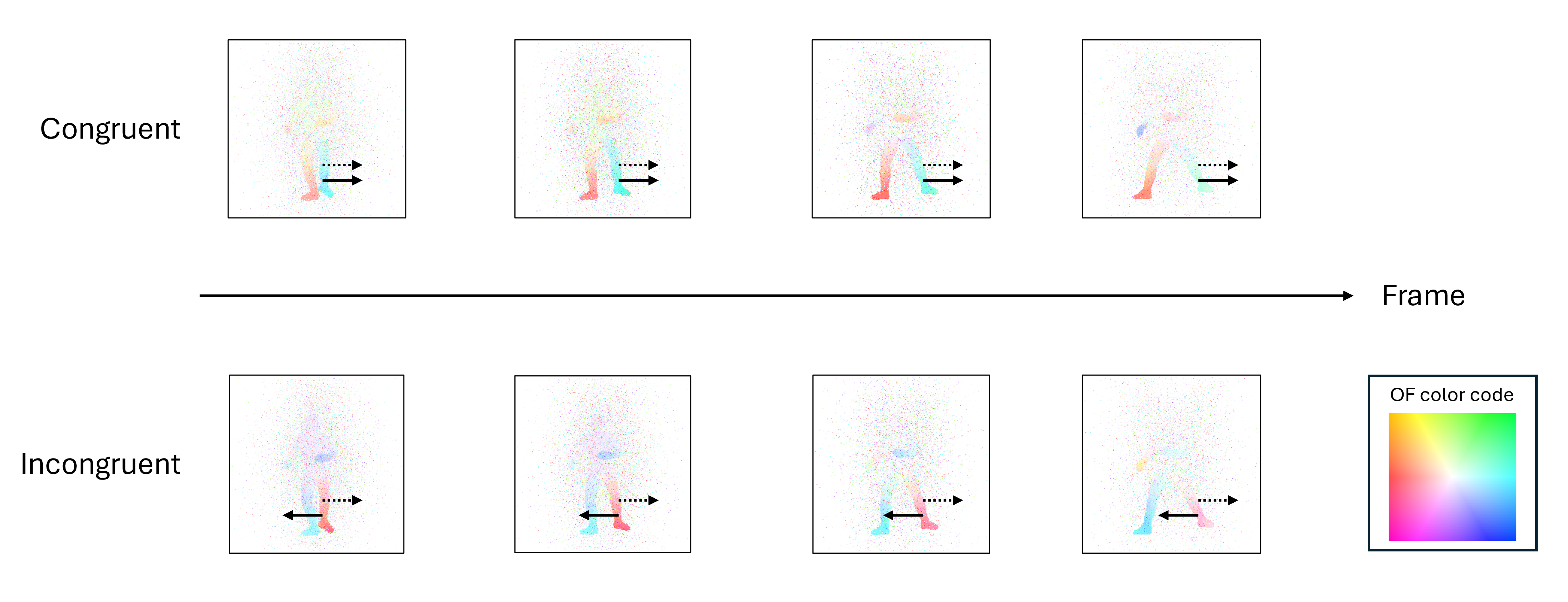

The congruence factor was controlled to modify the spatiotemporal dependence between the local motion vectors (solid arrow) and the displacement of motion field across time (dotted arrow). As shown in the plot, for both conditions, the motion field for one leg move to the right across time. For the congruent condition, each segment of local motion within the leg area has the same rightward velocity which matches the motion field displacement. However, for the incongruent condition, the local motion always has the opposite direction against the motion field displacement.

Supp. Figure 3. ROI response profiles for localizer conditions.

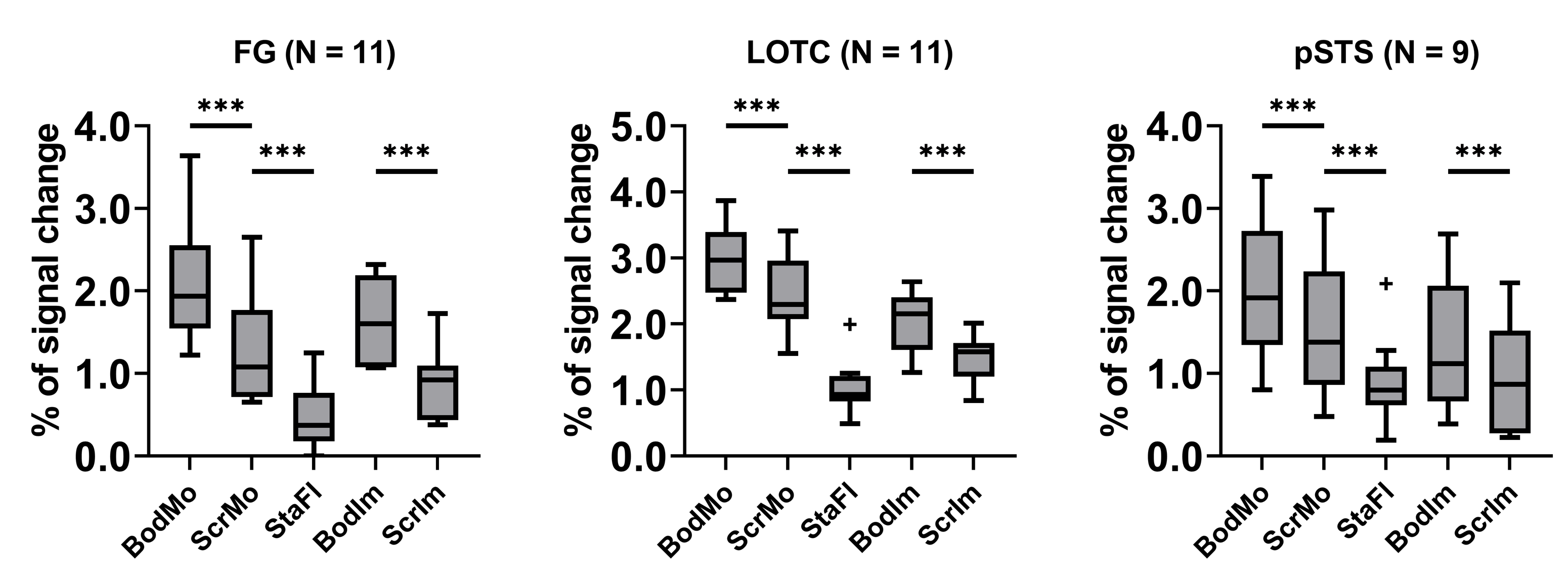

The boxplots show the beta values resulted from random effect ROI GLM on the localizer runs, with conditions of body motion (BodMo), scrambled dot motion (ScrMo), static flickering dots (StaFl), body image (BodIm), and scramble image (ScrIm). The whiskers (error bars) represent mininum / maximum values within 1.5 times the interquartile range from the lower or upper quartile and the scatters (+) are plotted for individual data beyond the upper and lower bounds. For all box plots, asterisks indicate the significant planned comparisons (Table 1) based on permutation-bootstrapping tests (*: p < 0.05; **: p < 0.01; ***: p < 0.001; two-tailed).

Supp. Figure 4. ROI response profiles for main experiment conditions.

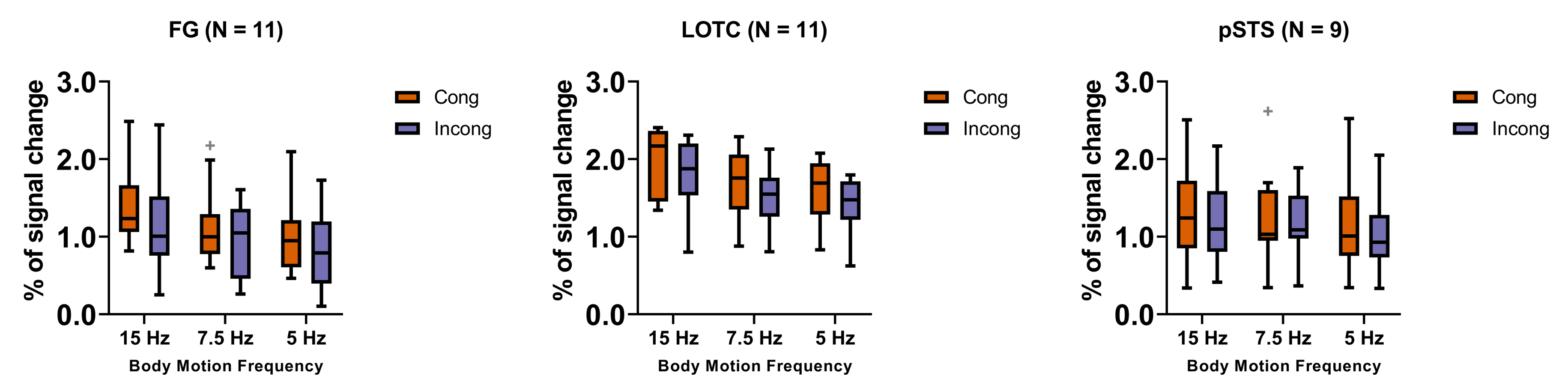

The boxplots show the beta values resulted from random effect ROI GLM on the main experiment runs, with whiskers (error bars) representing minimum / maximum values within 1.5 times the interquartile range from the lower or upper quartile and the scatters (+) plotted for individual data beyond the upper and lower bounds.

Supplementary Tables

### Supp. Table 1. Localizer experiment contrast design

|  | Body motion | Scrambled motion | Flickering dots | Body shape | Scrambled shape |
| --- | --- | --- | --- | --- | --- |
| Body motion selectivity | 1 | -1 | 0 | 0 | 0 |
| Dot motion selectivity | 0 | 1 | -1 | 0 | 0 |
| Body shape selectivity | 0 | 0 | 0 | 1 | -1 |

Supp. Table 2. Main experiment contrast design

|  | Cong / 15 Hz | Cong / 7.5 Hz | Cong / 5 Hz | Incong / 15 Hz | Incong / 7.5 Hz | Incong / 5 Hz |
| --- | --- | --- | --- | --- | --- | --- |
| Congruence effect | 1 | 1 | 1 | -1 | -1 | -1 |
| Frequency effect | 1 | 0 | -1 | 1 | 0 | -1 |
| Interaction | 1 | 0 | -1 | -1 | 0 | 1 |

Supp. Table 3. Details of subject-level ROIs.

| ROI | Subject | Hemisphere |  | MNI coordinates of center |  | voxeL number | t(df) |
| --- | --- | --- | --- | --- | --- | --- | --- |
|  |  |  | x | y | z |  |  |
| FG | 1 | L* | -44.31 | -36.44 | -20.32 | 79 | 5.69(1052) |
|  |  | R | 45.34 | -58.03 | -14.21 | 94 | 3.26(1052) |
|  | 2 | L | -47.53 | -65.22 | -15.21 | 391 | 7.42(1134) |
|  |  | R* | 45.5 | -56.34 | -17.04 | 706 | 8.02(1134) |
|  | 3 | R* | 45.16 | -54.71 | -16.94 | 419 | 5.30(1145) |
|  | 4 | L | -44.91 | -47.53 | -14.73 | 170 | 4.14(1117) |
|  |  | R* | 42.52 | -44.96 | -15.48 | 112 | 5.92(1117) |
|  | 5 | L | -41.51 | -48.43 | -12.32 | 84 | 4.21(1094) |
|  |  | R* | 43.92 | -44.67 | -16.57 | 96 | 5.04(1094) |
|  | 6 | L | -45.47 | -45.27 | -18.33 | 627 | 7.13(1085) |
|  |  | R* | 42.35 | -34.61 | -19.49 | 299 | 7.24(1085) |
|  | 7 | L* | -42.41 | -44.59 | -18.4 | 67 | 5.80(1104) |
|  | 8 | L* | -44.29 | -34.17 | -20.67 | 172 | 8.05(1129) |
|  |  | R | 46.18 | -53.01 | -15.31 | 953 | 7.15(1129) |
|  | 9 | L* | -44.43 | -52.98 | -17.89 | 67 | 6.34(1143) |
|  |  | R | 41.93 | -48.37 | -14.93 | 294 | 6.15(1143) |
|  | 10 | L* | -44.68 | -55.65 | -14.44 | 660 | 13.65(1123) |
|  |  | R | 42.49 | -47.69 | -19.47 | 774 | 8.41(1123) |
|  | 11 | L* | -43.67 | -42.42 | -18.21 | 174 | 6.60(1159) |
| LOTC | 1 | L | -54.84 | -64.35 | 5.02 | 933 | 6.18(1052) |
|  |  | R* | 49.4 | -64.13 | 9.27 | 1309 | 6.81(1052) |
|  | 2 | L | -46.18 | -73.27 | 10.41 | 3563 | 11.54(1134) |
|  |  | R* | 46.37 | -69.55 | 5.46 | 3337 | 11.83(1134) |
|  | 3 | L | -46.46 | -68.27 | 13.93 | 2362 | 8.55(1145) |
|  |  | R* | 46.87 | -66.63 | 12.89 | 2378 | 9.71(1145) |
|  | 4 | L | -46.08 | -75.83 | 9.91 | 2890 | 9.93(1117) |
|  |  | R* | 48.88 | -69.08 | 6.16 | 3835 | 11.05(1117) |
|  | 5 | L | -50.11 | -70.56 | 7.78 | 2174 | 7.78(1094) |
|  |  | R* | 48.5 | -68.48 | 8.42 | 2162 | 9.65(1094) |
|  | 6 | L | -51.03 | -71.7 | 11.42 | 2406 | 7.82(1085) |
|  |  | R* | 51.81 | -63.48 | 6.02 | 3103 | 8.07(1085) |
|  | 7 | L | -48.77 | -70.35 | 4.51 | 2499 | 9.36(1104) |
|  |  | R* | 50.78 | -62.83 | 3.14 | 3145 | 9.50(1104) |
|  | 8 | L* | -52.21 | -66.37 | 9.45 | 1958 | 8.88(1129) |
|  |  | R | 51.47 | -66.02 | 7.48 | 1152 | 7.47(1129) |
|  | 9 | L | -49.77 | -71.69 | 2.16 | 1405 | 7.31(1143) |
|  |  | R* | 47.64 | -66.81 | 3.83 | 3028 | 8.79(1143) |
|  | 10 | L | -48.49 | -69.16 | 9.02 | 4018 | 12.56(1123) |
|  |  | R* | 50.49 | -61.65 | 0.91 | 6034 | 15.42(1123) |
|  | 11 | L | -45.09 | -73.82 | 6.33 | 1328 | 8.85(1159) |
|  |  | R* | 45.88 | -66.28 | 1.36 | 2185 | 10.74(1159) |
| pSTS | 1 | R* | 54.22 | -46.72 | 11.26 | 382 | 4.57(1052) |
|  | 2 | L* | -55.82 | -46.1 | 15.06 | 447 | 5.30(1134) |
|  |  | R | 48.63 | -40.72 | 13.11 | 365 | 4.52(1134) |
|  | 3 | R* | 51.88 | -38.72 | 13.95 | 177 | 6.79(1145) |
|  | 6 | L | -51.07 | -46.99 | 11.43 | 353 | 5.79(1085) |
|  |  | R* | 53.97 | -40.15 | 13.33 | 1092 | 6.68(1085) |
|  | 7 | L* | -48.76 | -45.95 | 6.49 | 240 | 4.81(1104) |
|  |  | R | 53 | -37.26 | 8.61 | 212 | 4.69(1104) |
|  | 8 | L | -54.64 | -45.45 | 12.14 | 403 | 7.25(1129) |
|  |  | R* | 50.51 | -43.84 | 11.49 | 1975 | 9.18(1129) |
|  | 9 | L | -52.62 | -48.24 | 16.18 | 500 | 5.40(1143) |
|  |  | R* | 52.37 | -38.1 | 18.2 | 524 | 7.28(1143) |
|  | 10 | L* | -53.03 | -52.79 | 18.93 | 2466 | 11.91(1123) |
|  |  | R | 54.76 | -37.86 | 9.57 | 2493 | 9.47(1123) |
|  | 11 | L* | -55.83 | -39.08 | 9.08 | 557 | 7.52(1159) |
|  |  | R | 50.07 | -36.75 | 6.85 | 264 | 6.63(1159) |

Supp. Table 4. Voxelwise correlation between OF effects and localizer selectivity

| A. |  |  |  |  |
| --- | --- | --- | --- | --- |
|  |  |  | Flow motion effect |  |
|  |  | Congruence | Frequency | Interaction |
|  | Biological motion | r = 0.268, p < 0.001 | r = 0.283, p < 0.001 | r = 0.042, p = 0.612 |
| Localizer selectivity | Dot motion | r = 0.223, p = 0.004 | r = 0.349, p < 0.001 | r = 0.005, p = 0.956 |
|  | Body shape | r = -0.055, p = 0.469 | r = -0.168, p = 0.026 | r = -0.122, p = 0.104 |

| B. |  |  |  |  |
| --- | --- | --- | --- | --- |
|  |  |  | Flow motion effect |  |
|  |  | Congruence | Frequency | Interaction |
|  | Biological motion | r = 0.174, p = 0.005 | r = 0.204, p < 0.001 | r = 0.133, p = 0.035 |
| Localizer selectivity | Dot motion | r = 0.367, p < 0.001 | r = 0.598, p < 0.001 | r = -0.055, p = 0.561 |
|  | Body shape | r = -0.097, p = 0.043 | r = -0.096, p = 0.046 | r = -0.064, p = 0.184 |

| C. |  |  |  |  |
| --- | --- | --- | --- | --- |
|  |  |  | Flow motion effect |  |
|  |  | Congruence | Frequency | Interaction |
|  | Biological motion | r = 0.141, p = 0.076 | r = 0.237, p = 0.003 | r = 0.037, p = 0.648 |
| Localizer selectivity | Dot motion | r = 0.130, p = 0.113 | r = 0.253, p < 0.001 | r = -0.034, p = 0.668 |
|  | Body shape | r = -0.106, p = 0.104 | r = -0.010, p = 0.882 | r = -0.053, p = 0.424 |
